# Ultra-thin fluorocarbon foils optimise multiscale imaging of three-dimensional native and optically cleared specimens

**DOI:** 10.1101/533844

**Authors:** Katharina Hötte, Michael Koch, Lotta Hof, Marcel Tuppi, Till Moreth, Ernst H. K. Stelzer, Francesco Pampaloni

## Abstract

In three-dimensional light microscopy, the heterogeneity of the optical density in a specimen ultimately limits the achievable penetration depth and hence the three-dimensional resolution. The most direct approach to reduce aberrations, improve the contrast, and achieve an optimal resolution is minimizing the impact of changes of the refractive index along an optical path. Many light sheet fluorescence microscopes operate with a large chamber that contains an aqueous immersion medium and an inner specimen holder that contains the specimen embedded in a possibly entirely different non-aqueous medium. In order to minimize the impact of the specimen holder on the optical quality, we use multi-facetted cuvettes fabricated with vacuum-formed ultra-thin fluorocarbon (FEP) foils The ultra-thin FEP-foil cuvettes have a wall thickness of about 12 µm. They are resilient to fluidic exchanges, durable, mechanically stable and yet flexible.

We confirm the improved imaging performance of ultra-thin FEP-foil cuvettes with excellent quality images of whole organs, thick tissue sections and dense organoid clusters. The cuvettes outperform many other sample-mounting techniques in terms of full separation of the specimen from the immersion medium, compatibility with aqueous and organic clearing media, quick specimen mounting without hydrogel embedding, as well as their applicability for multiple-view imaging and automated segmentation. Additionally, we show that ultra-thin FEP foil cuvettes are suitable for seeding and growing organoids over a time period of at least ten days. The ultra-thin cuvettes allow the fixation and staining of the specimens inside the holder, preserving the delicate morphology of e.g. fragile, mono-layered three-dimensional organoids.

## Introduction

The light sheet fluorescence microscopy (LSFM) implementations of SPIM (single/selective plane illumination microscopy) and DSLM (digital scanned light sheet microscopy) ***(1)*** are optimal for the observation of large three-dimensional (3D) specimens with sizes ranging from micrometres (spheroids, organoids) to centimetres (biopsies, model organisms) ***(2–4)***. Ever since the invention of LSFM, agarose and other hydrogels have been used as transparent and biocompatible mounting media to immobilise specimens during imaging ***(5)***. While agarose-gel embedding is convenient for mounting drosophila and zebrafish embryos ***(6)***, it is unsuitable for most 3D cell cultures and especially inapplicable to mounting optically cleared specimens. In 3D cultures such as organoids, cells grow in soft hydrogels, e.g. Matrigel ***(7)***, which are composed of extracellular matrix (ECM) proteins such as collagen and laminin ***(8)***. Embedding 3D cultures into agarose gels deforms both the ECM hydrogels as well as the delicate multi-cellular structures growing inside. Attempts have been made to avoid specimen embedding, e.g. by depositing specimens into pre-cast “agarose beakers” ***(9, 10)***. However, the “agarose-beaker” mounting technique still presents many drawbacks including a time-consuming, laborious fabrication procedure and mechanical instability. Moreover, soluble components rapidly diffuse in and out of the agarose gel, implying that the specimens cannot be properly isolated from external contaminants of chemical or bacterial origin floating in the LSFM sample chamber ***(9)***.

Although three-dimensional specimens, ranging from organoids to organs are good models to investigate specific cellular processes in developmental biology and cancer research ***(11)***, it is often necessary to visualise the detailed architecture the whole organ or thick tissue section. In order to reduce strong light scattering effects caused by refractive index mismatches in three-dimensional, dense tissues, optical clearing solutions are used to homogenise the refractive indices across the whole sample. Two major families of optical clearing solutions are commonly used ***(12)***: (a) organic solvents and (b) water-based clearing solutions. Organic solvents are often chemically aggressive and can damage the objective lenses or the LSFM sample chamber. Thus, a common approach is using large glass or quartz cuvettes to contain the clearing medium in which the specimen is immersed ***(13)***. A second less corrosive medium is used to fill the LSFM sample chamber surrounding the cuvette. For this imaging approach, low magnification air objective lenses or macro lenses with low numerical apertures (NAs) are frequently used ***(14)*** yielding relatively low resolution and low image quality. Water-based clearing solutions are less corrosive compared to organic solvents, and can be combined with immersion objective lenses with higher NAs allowing for higher resolution and better image quality. Nevertheless, the handling of large volumes of optical clearing solutions can be challenging, e.g. due to cleaning issues or crystallisation of some components in the solution. In a previous work, we used thin square cross section glass capillaries with an inner wall width of 1 mm for mounting small organically cleared specimens. This allowed us to use immersion objective lenses in combination with less or non-corrosive media to fill the LSFM chamber in order to circumvent the drawbacks of the corrosive nature of organic clearing solutions ***(15, 16)***. Although we successfully applied this method to the *in toto* study of drug-treated spheroids, several disadvantages emerged. Not only does handling thin-walled glass capillaries bear potential risk of injury, but also attaching the capillaries to metal pins for placing them into the LSFM/DSLM sample chamber, and sealing the capillary/pin contact with Parafilm, wax, or similar proved to be a time-consuming and cumbersome procedure.

We introduced the use of fluorocarbon foil (fluorinated ethylene propylene, FEP) for LSFM in 2010, when we used FEP-cylinders to investigate the dynamics of microtubule asters in zebrafish embryos ***(11)***. FEP-cylinders filled with low concentrations of agarose or methylcellulose provided a more physiological confinement for developing embryos, and allowed for observation of microtubule dynamics unhindered by rigid and flat surfaces. This technique also proved to be suitable for long-term live imaging of zebrafish development ***(17)***.

Here, we describe how to fabricate and use ultra-thin FEP-foil cuvettes that eliminate the need for agarose/hydrogel embedding in LSFM imaging ***(18)***. The cuvettes are closed at the bottom end, which greatly simplifies handling compared to the previously used glass capillaries and FEP-cylinders. They allow fast and straightforward specimen mounting using forceps or pipettes to deposit the specimen into the cuvettes, and facilitate full containment of the internal mounting medium. The ultra-thin FEP-foil cuvettes are compatible with aqueous media, as well as with water-based and organic clearing solutions, which all differ in their refractive indices. Compared to agarose embedding, a key advantage of ultra-thin FEP-foil cuvettes is that the clearing medium of choice is confined within the cuvette, whereas the LSFM chamber itself can be filled with harmless and easy-to-handle index-matching media, such as 2,2′-Thiodiethanol (TDE), CUBIC2 or Iodixanol ***(19)***. A clear advantage of the ultra-thin FEP-cuvettes is minimising the effect of refractive index mismatch between FEP (n=1.34) and the surrounding mounting medium (water, n=1.33; TDE, n=1.41; CUBIC2, n=1.48; Iodixanol, n=1.75) by the reduced wall thickness of only a few microns (~12 µm). Our data show that these FEP-cuvette features allow high quality imaging of native (non-cleared) and optically cleared 3D specimens, such as hepatic and pancreatic organoids, thick murine kidney and brain tissue sections, as well as whole murine ovaries. The image quality allows for application of multi-view image reconstruction, and for subsequent application of automated quantitative image analysis pipelines.

## Material and methods

### FEP-foil

FEP-foil was purchased from Lohmann Technologies Ltd, UK (50 µm thickness, Batch No. GRN069662). Fluoroethylene propylene (FEP) is a thermoplastic fluorocarbon polymer composed of the monomers tetrafluoroethylene (CF_2_=CF_2_) and hexafluoropropylene (CF_2_=CF-CF_3_). FEP-films have outstanding properties for light microscopy. FEP-films are transparent with a refractive index of 1.341-1.347, close to the refractive index of water (1.333 at 20 °C). As most of the fluorocarbon plastics, FEP is chemically inert and resistant against organic solvents, acids and bases. Moreover, the material is autoclavable and biocompatible according to FDA (21CFR.177.1550) and EU (2002/72/EC) requirements (source: DuPont FEP-Fluorocarbon Film – Properties Bulletin).

### Fabrication of positive moulds for vacuum forming

We designed positive moulds of the cuvettes by using the free CAD software “123D Design” (Autodesk Inc., version 2.2.14) (**Figure 1a**, **Supplementary Figure 1a, b**). The 3D drawings were exported to stereolithography file format (.stl) for 3D printing. We 3D-printed the positive moulds by using the service of the company Shapeways (Eindhoven, NL, www.shapeways.com). The moulds were printed from UV cured acrylic polymer (commercially designed as “Ultra frosted detail”) in a Multijet Modeling (MJM) printing process. Two positive moulds, one with an array of square, and one with octagonal cross section pillars were 3D-printed (**Figure 1b**). The acrylic polymer has the required mechanical and thermal properties to withstand the vacuum forming process. The MJM printing process allows a detail resolution of 16 µm layer thickness, which is perfectly adequate for the preparation of the moulds. Before use, the positive moulds were inspected by stereomicroscopy and cleaned by immersion in an ultrasonic bath.

**Figure 1:**
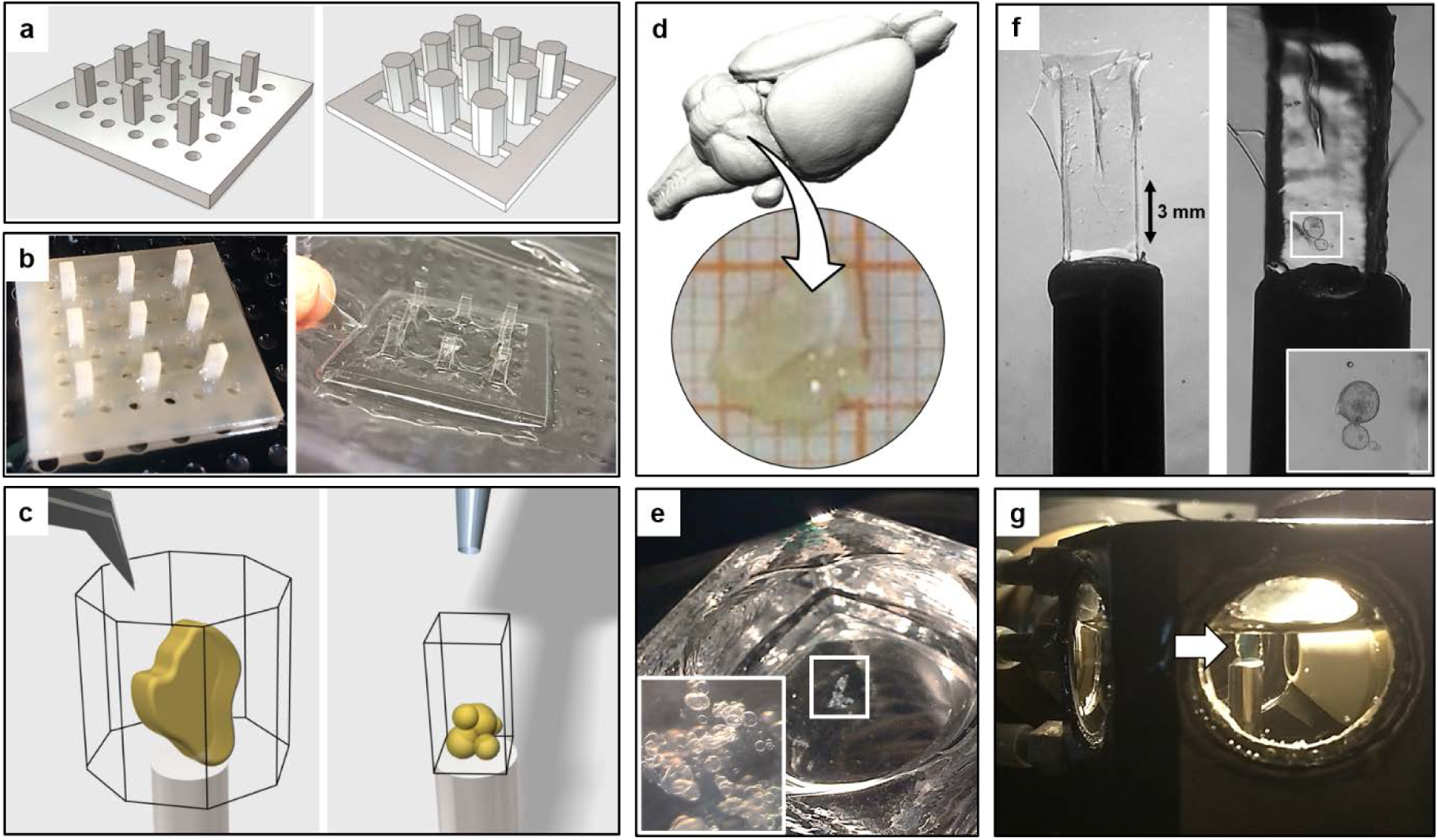
Fabrication of ultra-thin FEP-foil cuvettes and specimen preparation for LSFM. **(a)** CAD-derived drawings of positive moulds for arrays of 3×3 cuvettes. **(b)** Printed moulds (left) are used in the vacuum forming process of the ultra-thin FEP-foil cuvettes (right). **(c)** Sample preparation: an organ or a large tissue section (left) is positioned inside an ultra-thin FEP-foil cuvette with forceps. Smaller samples are deposited by pipetting (right). **(d)** A murine hippocampus section was isolated and optically cleared with CUBIC2. **(e)** Human hepatic organoids, partially embedded in Matrigel, are deposited in an embryo dish before being pipetted into the cuvette. **(f)** Ultra-thin FEP-foil cuvettes before (left) and after (right) human hepatic organoids are mounted. **(g)** Prior to image acquisition, ultra-thin FEP-foil cuvettes are glued to metal pins for inserting them into the mDSLM chamber.

### Cuvette fabrication with vacuum forming

In order to prepare a 3 x 3 array of vacuum-formed FEP-foil cuvettes, a 12 cm x 12 cm square patch of FEP-foil (Type A, general purpose, thickness 50 µm, DuPont de Nemours Int’l SA, Geneva, Switzerland) was clamped into the frame of a vacuum-forming machine (JT-18, Jin Tai Machining Company, Yuyao, PR of China). Once the heater raised the temperature close to the melting transition point of the FEP-foil (260-280°C), the 3D-printed positive mould was promptly placed onto the vacuum-forming machine, the vacuum suction was switched on, and the foil was quickly pressed onto the mould (**Figure 1b**, **Supplementary Figure 2**). Subsequently, the formed FEP-cuvette array was carefully removed from the mould with forceps, and cleaned with a detergent solution (1% Hellmanex-II in ultrapure water) in an ultrasonic bath for 10 minutes. The individual cuvettes were cut from the array with a scalpel. Next, individual cuvettes were glued onto stainless steel pins (diameter 3 mm, length 20 mm) using instant glue (**Figure 1c, f**).

**Figure 2:**
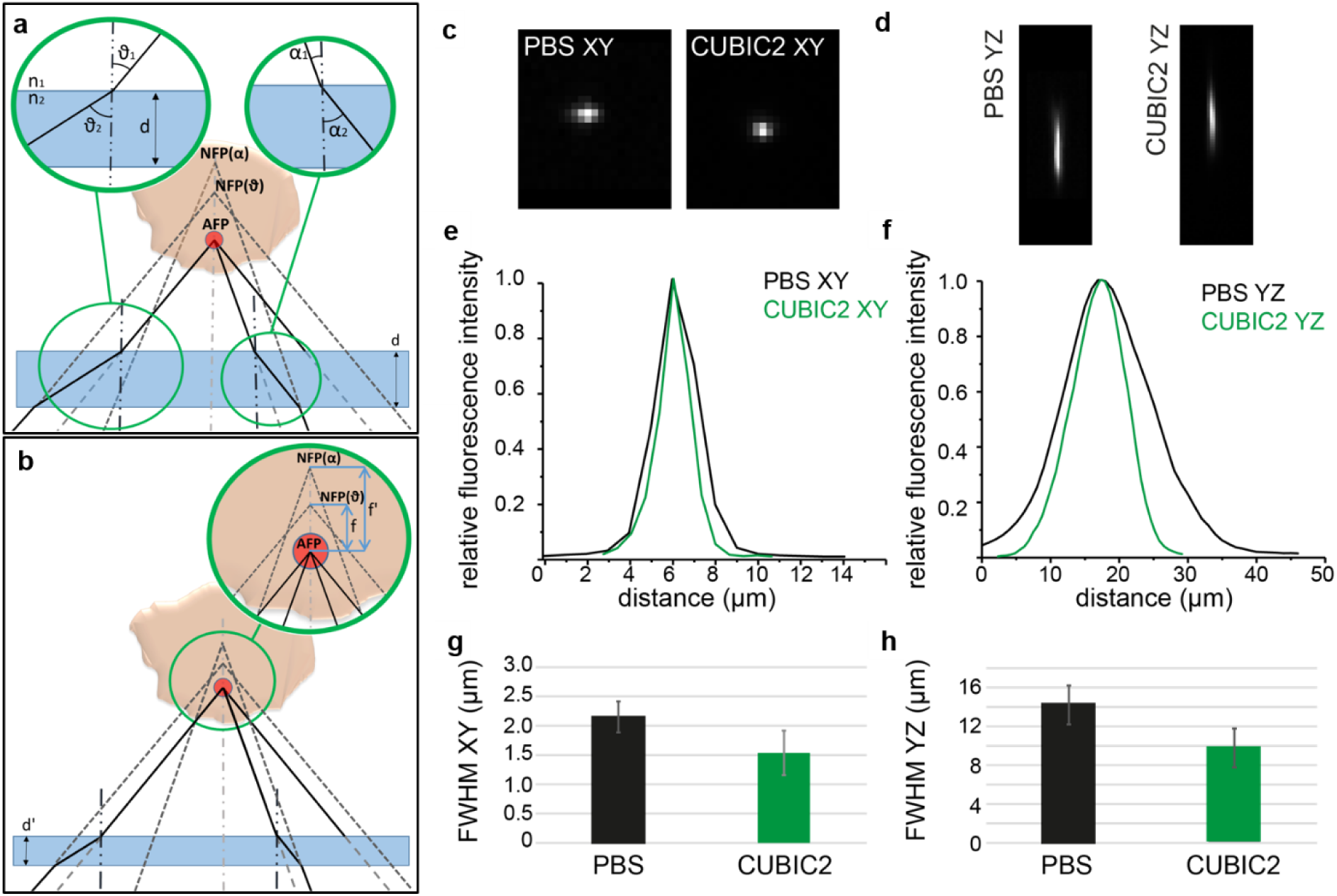
The reduced wall thickness of ultra-thin FEP-foil cuvettes minimises the spherical aberration caused by refractive index mismatches. **(a)** A cleared specimen is immersed in a medium with a refractive index *n*_1_ within an ultra-thin FEP-foil cuvette. The cuvette’s walls have a refractive index *n*_2_ with *n*_2_ > *n*_1_ and a thickness *d*. The cuvette is inserted in the LSFM chamber containing the clearing medium with the refractive index *n*_1_. Raytracing visualises the effects of refractive index mismatch, the thickness *d* and the angular dispersion on the spherical aberration. The light emitted by a fluorophore in the specimen (red dot) with a relatively large angle of incidence *v*_1_ with respect to the wall, is refracted at a relatively large angle *v*_2_ (left inset). Similarly, light at a smaller angle of incidence *a*_1_ is refracted at a smaller angle *a*_2_ (right inset). **(b)** The actual focal position (AFP) of the fluorophore (inset: *f*, *f*^′^) depends on the angle of incidence causing an aberration in the PSF. The relation between incidence and refraction angles is given by Snell’s law *n*_1_ *sin v*_1_ = *n*_2_ *sin v*_2_. By applying Snell’s law and simple trigonometric algebra, we can verify that the AFP is directly proportional to the thickness *d*, the refractive index ratio *n*_1_/*n*_2_, and the cosine of the angle of incidence *cos v*_1_: *f*~(*d*, *n*_1_/*n*_2_, *cos v*_1_). A smaller thickness *d*′ reduces the distance between the nominal focal position (NFP) and the AFP, thus minimising the spherical aberration. **(c - h)** Comparison of the Point-Spread Functions (PSFs) of fluorescent microspheres imaged within ultra-thin FEP-foil cuvettes in PBS and CUBIC2. **(c, d)** Images of the fluorescent microspheres in XY and YZ direction, respectively. XY-fluorescence intensity profiles of two representative microspheres (diameter 0.5 µm) at 488 nm in PBS and CUBIC2. Microscope: mDSLM. Objective lenses: Epiplan-Neofluar 2.5x/0.06 (excitation). N-Achroplan 10x/0.3 (detection). Excitation wavelength: 488 nm. Bandpass detection filter: 525/50 nm. **(e, f)** Fluorescence intensity profiles of two representative microspheres. **(g, h)** Comparison of the PSF’s full width half maximum (FWHM) values in PBS and CUBIC2 in XY and YZ direction. Error bars show the standard deviation (n=10). The FWHM was used as a relative measure for the PSF.

### Specimen preparation

#### Murine brain

We obtained adult murine brains from a transgenic strain expressing GFP under the control of the Thy-1 promoter (kindly provided by Dr. Walter Volknandt, Goethe Universität Frankfurt, Faculty of Neurobiology and Biosciences). The brains were fixed in 4% PFA in PBS at 4°C overnight. Prior to clearing, the brains were washed three times with PBS for 15 minutes, and the hemispheres were separated. Next, the hemispheres were cut into smaller blocks, with a size of approximately 5 mm x 5 mm x 1.5 mm (**Figure 1d**). The blocks were incubated with CUBIC2 clearing solution in a reaction tube overnight. Before imaging, the CUBIC2 solution was refreshed, and the blocks were inserted into 5 mm-diameter octagonal FEP-cuvettes using forceps (**Figure 1c**).

#### Murine kidney

The kidney was fixed in 4% PFA in PBS at 4°C overnight. Prior to clearing, the kidney was washed three times with PBS for 15 minutes, and cut in small blocks.

#### Murine ovary

Explanted and *ex vivo* cultured ovaries, expressing GFP-cKit specifically in all oocytes, were treated as described ***(20, 21)***. Staining was performed as described by Smyrek *et al.* with slight modifications ***(16)***. Briefly, ovaries were harvested from eight-day-old (P8) female GFP-cKit mice and fixed in 4% PFA in PBS overnight. The ovaries were permeabilised with 0.3% Triton X-100 in PBS for 30 minutes at room temperature in a 96-well flat-bottom plate (Greiner), while shaking at 450 rpm. Next, the ovaries were treated with blocking buffer (0.3% Triton X-100, 0.05% Tween-20, 0.1% BSA and 10% donkey serum in PBS) for 2 hours at room temperature. After blocking, the ovaries were incubated with DAPI (Thermo Fisher; 1 μg/ml in blocking buffer) at 37 °C in a humidified incubator in the dark for 24 hours. Afterwards, the ovaries were washed three times with PBS in the dark for 20 minutes, and were kept in PBS at 4°C, protected from light in a humidified incubator. The ovaries were washed four times in CUBIC2 solution, before being transferred with forceps into the octagonal FEP-cuvettes containing CUBIC2 solution (**Figure 3c-d**). In this step, it is important that no air bubbles enter the cuvette, to avoid floating of the ovaries.

**Figure 3:**
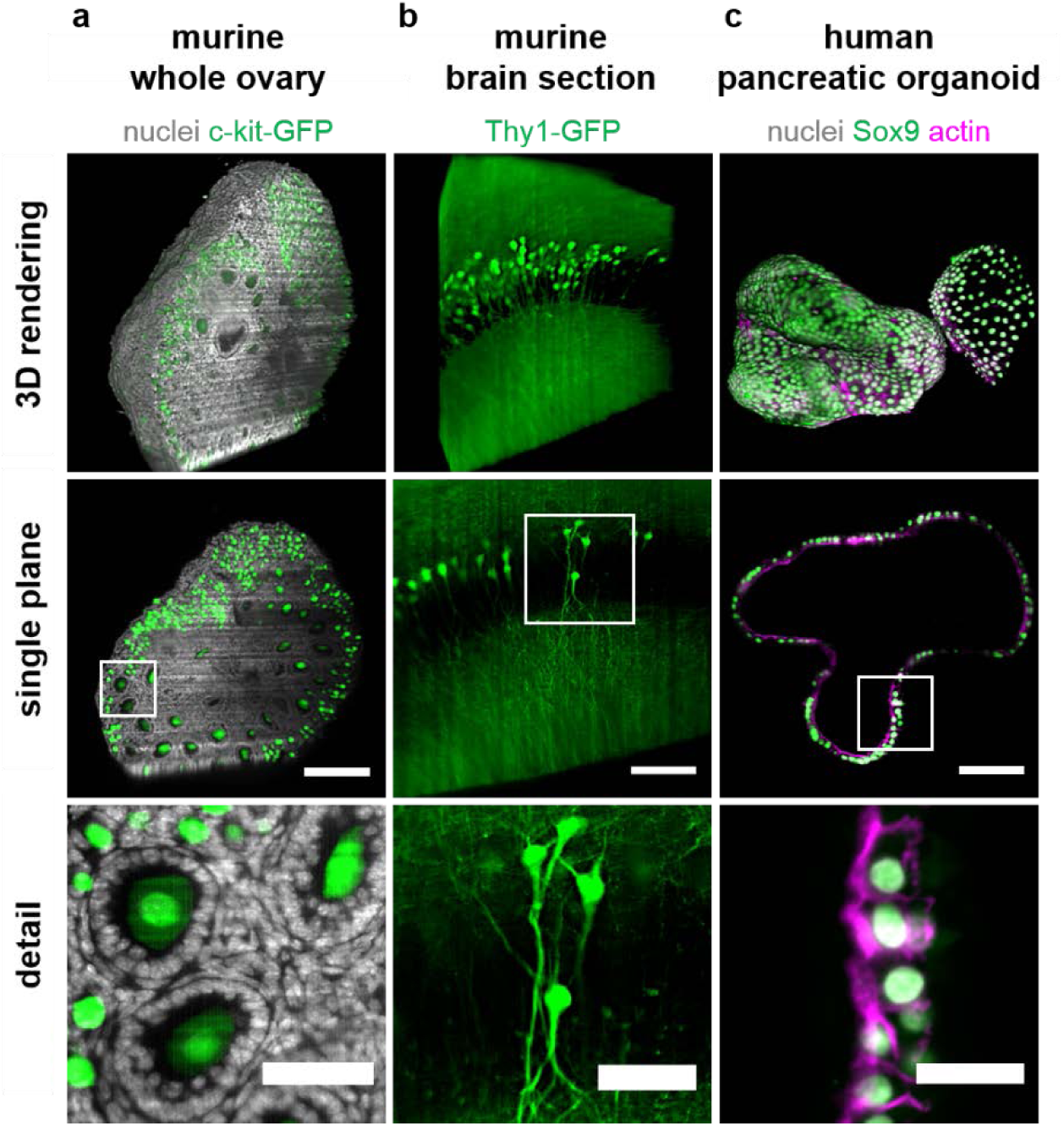
Images of specimens with heterogeneous optical densities mounted into ultra-thin FEP-foil cuvettes provide high resolution at the subcellular level. Shown are 3D renderings, single central planes and detail views of **(a)** a whole murine ovary of an 8-day old transgenic p-18 GFP-c-Kit mouse, in which all oocytes express GFP-c-Kit (green), stained with DAPI (grey), optically cleared with CUBIC2, showing primordial and primary oocytes, as well as the nuclei of surrounding cells, e.g. granulosa cells, **(b)** a thick (1.5 mm) murine brain section of a mouse expressing GFP under the control of the Thy-1 promoter, showing GFP-expressing cells in the granular layer within the dentate gyrus, optically cleared with CUBIC2, and **(c)** a native human pancreatic organoid immuno-stained against the pancreatic progenitor cell marker Sox9 (green), stained with phalloidin (magenta) and DAPI (grey). All samples were imaged with LSFM in ultra-thin FEP-foil cuvettes. Microscope: mDSLM. Objective lenses: Epiplan-Neofluar 2.5x/0.06 (excitation). N-Achroplan 10x/0.3 (detection). Excitation wavelength: 488 nm. Bandpass detection filter: 525/50 nm. Scale bar: 100 µm, detail view scale bar: 50 µm.

#### Human and murine organoids

Human adult hepatic organoids were obtained from Luc JW van der Laan (Erasmus MC – University Medical Center Rotterdam, NL). Hepatic organoids were cultured in expansion medium for 7 days as previously described ***(22)***. Murine and human adult pancreatic organoids were obtained from Meritxell Huch (Gurdon Institute, Cambridge, UK). Pancreatic organoids were cultured as previously described ***(7)*** with adjustments for human pancreatic organoids (Meritxell Huch, personal communication, unpublished data). Isolation of organoids from the embedding matrix (human hepatic organoids: Matrigel, Corning; human pancreatic organoids: Cultrex BME2, Amsbio) for fixation and whole-mount staining was performed with slight modifications of protocols published by Broutier *et al*. ***(23, 24)***. Briefly, organoids were extracted from the matrix by washing three times with ice-cold 0.1% BSA in PBS. Organoids were fixed with 2% PFA in PBS on ice for 30 min. Whole-mount staining of the specimens was done using an adapted version of a protocol previously described by Smyrek *et al. **(16)***. In short, organoids were incubated with DAPI (Thermo Fisher; 1 μg/ml in PBS). Human pancreatic organoids were additionally immuno-stained against Sox9 (primary antibody: Merck Millipore, AB5535; secondary antibody: goat anti-rabbit IgG (H+L) Alexa Fluor 488, Thermo Fisher, A11008), and labelled with Alexa Fluor 568 Phalloidin (Thermo Fisher, 1:200). The stained specimens were washed three times and stored in PBS at 4°C. Hepatic organoid clusters, still partially embedded in Matrigel, were gently transferred from an embryo dish (**Figure 1e**) into square cross section FEP-cuvettes attached to stainless steel pins (**Figure 1c, f**) using a pipette with a 100 µl tip cut at the end in order to avoid damaging of the specimen due to shear forces. The cuvette was pre-filled with PBS. PBS was used as both mounting (in the cuvette) and imaging medium (in the DSLM chamber, **Figure 1g**). The same mounting procedure was used for single human pancreatic organoids. For nuclei segmentation, murine organoids were grown within ultra-thin FEP-foil cuvettes for three days (**Supplementary Figure 6**). Prior to organoid seeding, the FEP-cuvettes were sterilised in 75% Ethanol for at least 3 hours, washed thoroughly with PBS and dried. They were then filled with 20 µl of Matrigel containing organoid fragments, placed into a 48-well plate, and fully covered with expansion medium. After three days of cultivation, the FEP-cuvettes containing the organoids embedded in Matrigel were briefly washed with PBS, and fixed with 4% PFA in PBS supplemented with 1% glutaraldehyde (Electron Microscopy Sciences, 16020) for 30 minutes at 4°C. After two short washing steps with PBS, up to three FEP-cuvettes were pooled in a 1.5 ml tube and stained with DAPI (Thermo Fisher; 1 μg/ml in PBS) overnight. For imaging, the filled FEP-cuvettes were attached to stainless steel pins using instant glue. PBS was used as imaging medium to fill the DSLM chamber.

### Optical clearing

#### Whole murine ovaries and brain sections

For clearing of whole murine ovaries ***(20)*** and large tissue sections of murine brain, we used CUBIC2 clearing solution ***(25)***. Briefly, 50% w/v sucrose, 25% w/v urea and 15% w/v deionized water were mixed and stirred at 60°C. After all components were dissolved, 10% w/v 2,2′,2′’-nitrilotriethanol was added and stirred at room temperature. The clearing solution was de-gassed by placing it into a desiccator for about 20 minutes. Finally, the refractive index was determined (n = 1.49). The specimens were immersed in CUBIC2 in a reaction tube overnight.

#### Murine kidney

Optical clearing of murine kidney was performed with ethyl cinnamate (ECi, n = 1.58) ***(26)***. The specimen was dehydrated with increasing concentrations of ethanol in ultrapure water. After the last dehydration step with 100% ethanol, the specimen was immersed in ECi in a reaction tube overnight.

### Cleaning procedure for re-using the ultra-thin FEP-foil cuvettes

The imaged specimen was removed and the cuvettes were transferred into a 15 ml reaction tube. Next, the cuvettes were washed in a 1% solution of Hellmanex-II in ultrapure water at room temperature on a rotator overnight. After washing, the Hellmanex-II (Hellma Analytics, Müllheim, DE) solution was discarded and the cuvettes were rinsed twice with ultrapure water for at least one hour on the rotator. After drying at 60°C for at least one hour, the cuvettes can be re-used. Slight deformation of the cuvette’s walls caused by the washing procedure can be corrected by gently pressing with tweezers.

### LSFM imaging

The FEP-cuvettes were glued onto stainless steel holders (**Figure 1f**), and images were acquired with a custom-built monolithic digital-scanned light sheet-based fluorescence microscope (mDSLM) ***(6)***, which features a motorized xyzϑ-stage placed below the specimen chamber (**Figure 1g**). The microscope was equipped with an Epiplan-Neofluar 2.5x/0.06 illumination objective (Carl Zeiss) and an N-Achroplan 10x/0.3 detection objective (Carl Zeiss), and a Clara CCD camera (ANDOR Technology, Ireland). Laser and bandpass filter sets: 561 nm, 607/70; 488 nm; 525/50, 405 nm, 447/50.

### Measurement of the point spread function (PSF)

The fluorescence intensities of single TetraSpeck microspheres (Thermo Fisher) were used as a measure for the relative axial (yz) and lateral (xy) fluorescence intensity distribution in PBS and CUBIC2 microscopy immersion media, respectively. Microspheres were embedded with low-melting point agarose into square cross sections FEP-cuvettes. We used a custom-built mDSLM equipped with an Epiplan-Neofluar 2.5x/0.06 illumination objective (Carl Zeiss) and an N-Achroplan 10x/0.3 detection objective (Carl Zeiss). A Clara CCD camera (ANDOR Technology, Ireland) was used for image acquisition. The microspheres were excited at 488 nm. A bandpass filter centred at 525/50 was used to record the fluorescence emitted by the beads. Images of microbeads close to centre of the agarose column were recorded and analysed with Fiji (ImageJ version 1.51d, Java version 1.6.0_24). Line intensity profiles were measured by using the line selection tool of Fiji and the Analyze/Plot Profile tool. The image of each bead was normalized by dividing by the maximum pixel value in each image. The intensity profiles of 10 fluorescent microbeads along the XY- and YZ-direction were analysed for each condition. After a Gaussian fitting of the lateral and axial distributions, the full width half maxima (FWHM) were determined as a measure for the resolution ***(27)***.

### Image processing

Raw image stacks were pre-processed with Fiji (ImageJ version 1.51d, Java version 1.6.0_24) ***(28)***. The stacks were cropped to the region of interest. The background intensity was subtracted from every slice using the function Subtract Background (Fiji, ball radius of 60 pixels). Next, the stacks were resliced with a factor of 4 along the z-axis. 3D maximum intensity projections were generated with Fiji (projection method: brightest point, 360° rotation angle, angular increments were set at 1° or 10°). The multi-view registration and fusion of the organoid stack displayed in Figure 4 was performed with Huygens Essential SPIM/Light Sheet Fusion & Deconvolution Wizard (Scientific Volume Imaging, NL) ***(29)***. The volume rendering of the ovary in Figure 3 was performed with the Fiji plugin 3D viewer ***(30)***. Stripe filtering and deconvolution of the murine kidney dataset shown in Supplementary Figure 4 were conducted with the Fiji plugins Parallel Iterative Deconvolution 3D v1.12 ***(31)*** and with the VSNR V2 Variational Stationary Noise Remover ***(32)***, respectively. In the VSNR plugin, a Gabor filter with a 90° angle, sigma_x = 3, sigma_y = 100, Lambda = 0, and a noise level = 0.3 were selected. The stack was then processed in Mode 2D.

**Figure 4:**
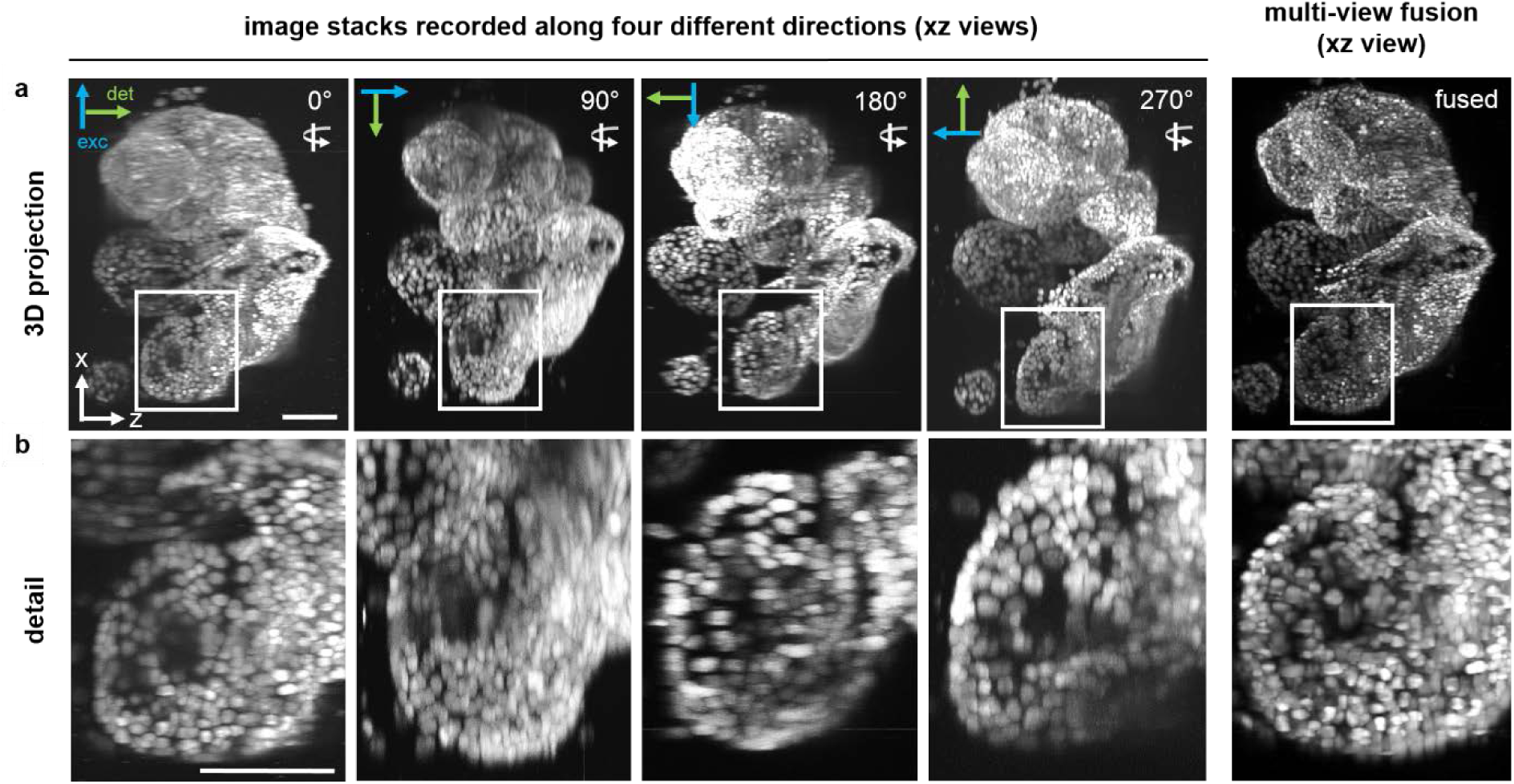
Projections through image stacks of human hepatic organoids recorded along four directions for multi-view fusion. A cluster of human hepatic organoids was carefully inserted into an ultra-thin square cross section FEP-foil cuvette. Four stacks consisting of 301 images each were recorded with a DSLM along four different directions, i.e. rotation angles around the Y-axis of 90°, 180° and 270°. Shown are four 3D maximum intensity XZ-projections that provide views of the specimen “from above”, i.e. along the Y-axis. The projections for each rotation angle are aligned to show the specimen along a comparable orientation. Arrows indicate the orientations of the light paths of excitation (blue) and detection (green) for each angle. **(a)** Overviews of the entire organoids cluster. **(b)** Detailed views of one organoid in the cluster. Staining: cell nuclei (DAPI). Microscope: mDSLM. Objective lenses: Epiplan-Neofluar 2.5x/0.06 (excitation). N-Achroplan 10x/0.3 (detection). Excitation wavelength: 405 nm. Bandpass detection filter: 447/25 nm. Scale bars: 100 µm.

### Automated nuclei segmentation

Nuclei segmentation was performed as previously described in Schmitz et al. 2017. Raw image stacks of the nuclei channel were pre-processed in Fiji (ImageJ version 1.51h, Java version 1.8.0_66). The raw image stacks were cropped to the region of interest. Cell nuclei were extracted by means of an automated three-dimensional segmentation pipeline implemented in Wolfram Mathematica (version 10.3). In brief, pre-processed image stacks were rescaled by a factor of 4 along z-axis to obtain isotropic voxels. Hence, in the resized image, the voxels are isotropic with a pitch of 0.654 µm. The images were then deconvolved with a median filter range of three pixels. Cell nuclei segmentation was achieved by local thresholding and a marker-controlled immersion-based watershed algorithm. The local threshold corresponds to the mean of the local intensity distribution in a range of ten voxels. Marker points for the watershed algorithm were detected by a multiscale Laplacian of Gaussian seed detector with a minimal range of eight and a maximum range of twelve voxels ***(33, 34)***. Marker points were increased by morphological dilation with a spherical structuring element and a radius of three voxel. Using the marker points as starting points, the final segmentation result was obtained by an immersion-based watershed algorithm ***(35)***. Using Edelsbrunner’s algorithm ***(36)***, the cell nuclei centroids were used to compute an alpha shape with the alpha parameter set to 240 voxels. From the alpha shape, the approximate spheroid volume could be readily obtained. A cell graph ***(37, 38)*** was constructed with an edge distance threshold of 60 voxels capturing the spatial arrangement of cells and the local cell density within an organoid. The maximal outlier distance was 55 voxels. Features of all cell nuclei that had a volume greater than 500 and smaller than 18000 voxels, and global features of the spheroid were stored in tabular format.

## Results

### Fabrication of ultra-thin FEP-foil cuvettes can be easily adapted to specimen type and size

For the fabrication of ultra-thin FEP-foil cuvettes, we sketched two positive moulds of the cuvettes for 3D printing (**Figure 1a**). One mould was designed as a square cross section array to produce cuvettes that are suitable for embedding smaller specimens in a size range of micrometres to millimetres. The other mould was designed to produce cuvettes with an octagonal cross section, allowing the mounting of larger specimens in a size range of several millimetres to centimetres. The moulds were 3D-printed from an UV-cured acrylic polymer (commercially designed as “ultra frosted detail”) in a Multijet Modeling (MJM) printing process (**Figure 1b**, left). The acrylic polymer possesses the required mechanical and thermal properties to withstand the vacuum forming process during cuvette fabrication. The MJM printing process allows a detail resolution of a 16 µm layer thickness, which is perfectly adequate to achieve the detail precision needed for the production of miniaturized moulds. The ultra-thin FEP-foil cuvettes were produced using a vacuum-forming machine. In brief, the FEP-foil was heated to its glass transition point (260-280°C) before being stretched over the acrylic moulds. The fabrication process is described in more detail in the methods section. On one hand, this process facilitates the moulding of a durable but flexible shape. On the other hand, the expansion of the FEP-foil by a factor of about 4 allows the formation of ultra-thin FEP-foil cuvettes with a wall thickness of about 12 µm (**Supplementary Figure 2**). Single cuvettes are cut from the array (**Figure 1b**, right) with scissors, and are glued onto a metal pins (sample holders) before depositing the specimen together with mounting medium into the cuvettes (**Figure 1f**). As these ultra-thin FEP-foil cuvettes are durable but flexible, small and large specimens such as dense organoid clusters (**Figure 1e**) and thick murine brain sections (**Figure 1d**) can be easily deposited and positioned inside the cuvette using forceps or pipettes (**Figure 1c**). Finally, the FEP-cuvette/metal pin sample holders are placed into the LSFM sample chamber filled with imaging medium (**Figure 1g**).

### Optical properties of ultra-thin FEP-foil cuvettes outperform commonly used materials for specimen mounting in LSFM

**Figure 2a, b** illustrate the impact of the foil thickness on the refractive index mismatch. The spherical aberration caused by the refractive index mismatch depends (also) on the foil thickness. The stretching of the foil introduced by the vacuum-forming process reduces the wall thickness, minimising the image deterioration caused by index mismatches. To characterise the imaging properties of the ultra-thin FEP-foil cuvettes, we compared the point spread functions (PSFs) of fluorescent microspheres inside the FEP-cuvettes with a DSLM setup using different water-based imaging media. To this purpose, the microspheres were embedded in PBS/agarose gel inside the cuvettes, which were subsequently immersed in the DSLM sample chamber filled with either PBS or CUBIC2 ***(39)***. Images of the lateral (XY) and axial (YZ) PSFs of the fluorescent microspheres in PBS and CUBIC2 are shown in **Figure 2c, d**, respectively. **Figure 2e, f** display the corresponding lateral and axial extensions of the PSFs in PBS and CUBIC2. Despite the refractive index matching being sub-optimal for the CUBIC2 setup, in which the emitted light has to pass through the PBS/agarose gel inside the cuvette, FEP (n=1.34), and CUBIC2 (n=1.48), the full width half maximum (FWHM) values of the lateral and axial PSFs in CUBIC2 are significantly smaller (30%) than the ones in PBS (**Figure 2g, h** and **Supplementary Table 1**).

### Ultra-thin FEP-foil cuvettes are applicable to *in toto* imaging of a wide variety of specimens

In order to test the applicability of the ultra-thin FEP-foil cuvettes we imaged with light sheet fluorescence microscopy a variety of biologically diverse specimens applying different sample mounting techniques. The specimens included CUBIC2-optically cleared whole murine ovaries, thick murine brain sections, ethyl cinnamate (ECi)-cleared murine kidney sections, and native (non-cleared) human pancreatic organoids (**Figure 3**). Individual whole murine ovaries expressing GFP-c-kit were DAPI-stained, cleared with CUBIC2, and placed into the square cross section FEP-cuvettes for imaging (see **Movie 1**). With a 10x/0.3 objective lens, it was possible to image the entire densely-packed organ without need for post-acquisitional image stitching. At the same time, we could easily distinguish between the stromal cells, the boundary of the *teca*, the cell layers in the *granulosa*, and the cytoplasm of the oocytes (**Figure 3a**, detail view). **Supplementary Figure 3** shows 25 optical sections through the whole ovary of an 8 days-old (P8) mouse. We achieved equally good results in the microscopy of thick murine brain sections in which GFP was expressed under the control of the Thy-1 promoter (see **Movie 2**). **Figure 3b** (single plane, detail view) shows an optical section of the granular layer in the dentate gyrus of the murine hippocampus. The cleared brain slice with a size of approximately 5 mm x 5 mm was inserted into an octagonal cross section FEP-cuvette. A suitable region of interest (ROI) was selected and imaged with a DSLM. A maximum intensity projection of a Z-stack composed of 101 slices, a three-dimensional surface rendering, and a detail view show that the image quality is sufficient for morphological investigations of lipid-rich tissues. The image shows a single ROI, and no subsequent image stitching was performed. The results of ovary and brain imaging demonstrate that the combination of CUBIC2-clearing and the ultra-thin FEP-foil cuvettes allows an *in toto* analysis of mammalian organs and thick tissue slices. In order to determine whether ultra-thin FEP-foil cuvettes also provide good image quality in combination with organic clearing agents with refractive index n~1.5, a murine kidney section was cleared with ethyl cinnamate (ECi) and imaged by recording the auto-fluorescence signal of the tissue excited at 488 nm (see **Supplementary Movie 1**). The image quality was excellent, despite the large refractive index mismatch between the FEP-foil (n=1.34) and ECi (n=1.56) (**Supplementary Figure 4**). We further tested the performance of the ultra-thin FEP-foil cuvettes in combination with immuno-labelled organoids (see **Movie 3**). Therefore, human pancreatic organoids were DAPI-stained, immuno-labelled against the pancreatic progenitor marker progenitor marker Sox9 ***(40)***, and phalloidin-stained to visualise the actin skeleton (**Figure 3c**). The close-up provides a detailed overview of the spatial distribution of the actin cytoskeleton, which is most prominent at the membranes facing the organoid lumen. In summary, these data suggest that the ultra-thin FEP-foil cuvettes are applicable to a wide variety of biologically diverse specimens, including native (non-cleared) and optically cleared specimens, and allow the microscopic assessment of subcellular structures in large specimens.

### Ultra-thin FEP-foil cuvettes are suitable for multi-view imaging and three-dimensional multi-view reconstruction

For delicate multi-cellular structures such as organoids, comprising a cell-monolayer and hollow lumen, it is often not possible to perform optical clearing to reduce light scattering and improve image quality. Attempts to replace the water in the specimen with an optical clearing agent often leads to damage of the original architecture and collapse. For these kinds of specimens, the possibility to easily perform multi-view imaging is essential in order to achieve high-quality and isotropic 3D image stacks, suitable for quantitative image-based analysis. To test the applicability of the ultra-thin FEP-foil cuvettes for multi-view imaging of complex 3D cell cultures, we imaged a DAPI-stained dense cluster of human hepatic organoids still partially embedded in Matrigel at four rotation angles (0°, 90°, 180°, 270°) in a square cross section FEP-cuvette (**Figure 4**). The cluster of cyst-like organoids growing in very close proximity to each other (**Supplementary Figure 5**), together with the residual Matrigel, represent a complex and highly light-scattering specimen. The small size of the square cross section FEP-cuvette (side length: 2 mm) allows for an unimpeded 360° rotation of the specimen in the mDSLM sample chamber. 3D maximum intensity XZ-projections of the image stacks recorded along four different directions are presented in **Figure 4a**. Each projection was rotated for alignment and comparability with the 0°-projection. The detail views of the 0°-, 90°-, 180°-, and 270°-projections of the organoid cluster (**Figure 4b**) reveal optical distortions due to light scattering that commonly occur when imaging complex and highly scattering samples. Multi-view imaging and multi-view fusion significantly reduce the optical distortions, and provide more detailed information on the subcellular level as shown in the detail view of the fused data set (**Figure 4b**, also see **Movie 4**). These data provide evidence of the improvement of the image quality resulting from multi-view imaging with ultra-thin FEP-foil cuvettes and multi-view fusion of native (non-cleared) specimens.

**Figure 5:**
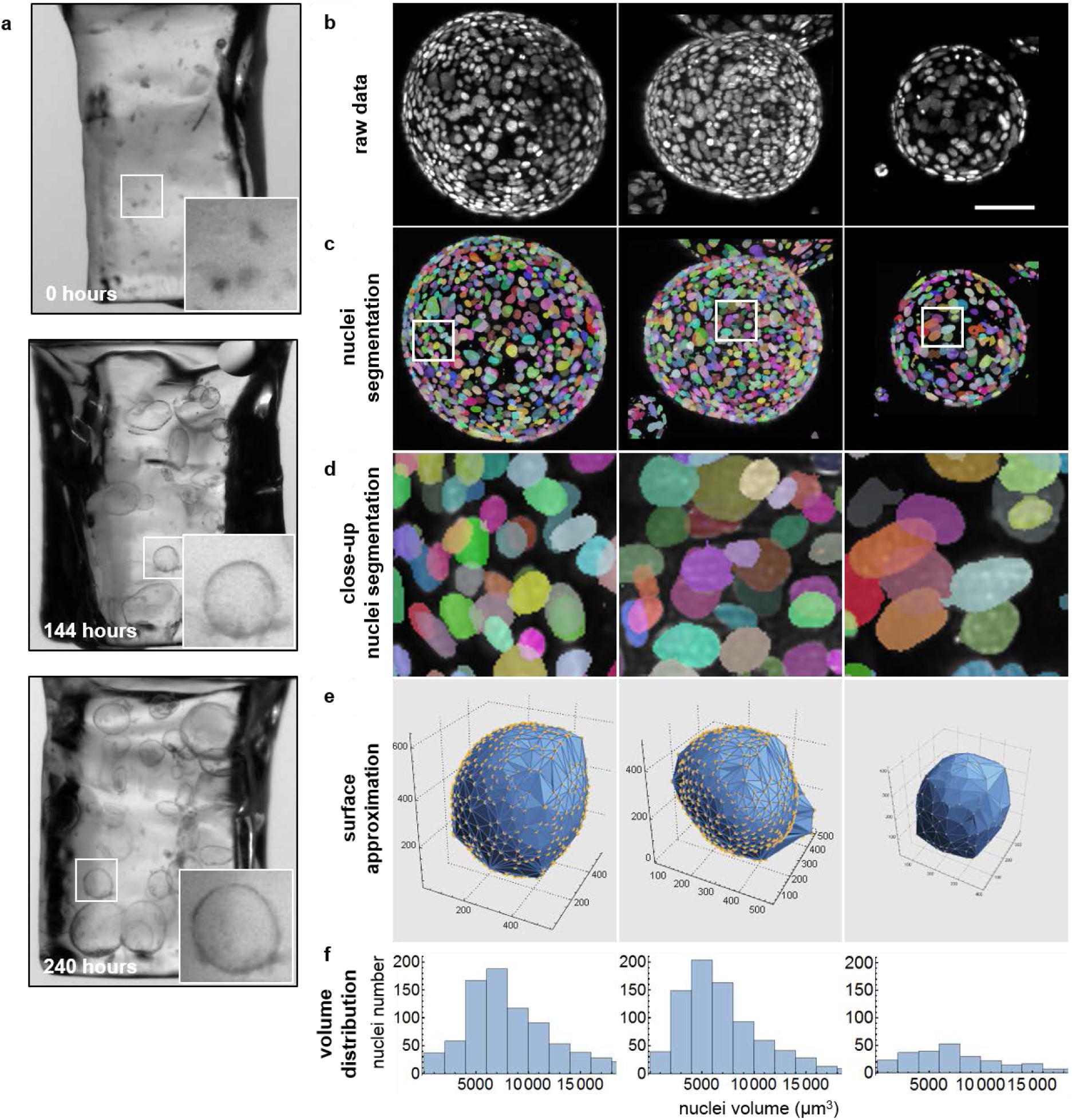
Long-term culture of murine pancreatic organoids inside ultra-thin FEP-foil cuvettes for maintain unimpaired morphology during fixation and labelling as prerequisite for three-dimensional image segmentation. **(a)** Organoids were directly seeded into ultra-thin FEP-foil cuvettes and cultured over a period of ten days (240 hours). The insets show healthy pancreatic organoids 0, 144 and 240 hours after seeding. The organoids remain in the same cuvette during fixation, labelling and imaging. **(b)** Maximum intensity XZ-projections of raw image stacks of three different pancreatic organoids after fixation and DAPI staining inside the ultra-thin FEP-foil cuvette. Microscope: mDSLM. Objective lenses: Epiplan-Neofluar 2.5x/0.06 (excitation). N-Achroplan 10x/0.3 (detection). Excitation wavelength: 405 nm. Bandpass detection filter: 447/25 nm. Scale bar: 100 µm. **(c)** High image quality ensures an optimal separation amongst the labeled nuclei, which is essential for semi-automated nuclei segmentation. Different colors indicate individual nuclei. **(d)** Close-ups of segmented nuclei. **(e)** The global morphology of the organoids is shown in surface approximations based on alpha shapes. **(f)** Histograms show the nuclei volume distribution within the three different organoids.

### Ultra-thin FEP-foil cuvettes are suitable for long-term cultivation of organoids and facilitate high quality imaging for quantitative image-based analysis

Image-based analyses require a good signal to noise ratio and a sufficient resolution of subcellular structures. We aimed to culture organoids embedded in matrix directly inside the ultra-thin FEP-foil cuvettes in order to preserve their morphology during fixation and staining, and assess the image quality in the subsequent imaging process (**Supplementary Figure 6**). After seeding murine pancreatic organoid fragments into the FEP-cuvettes, the filled cuvettes were placed in a culture dish and were covered with expansion medium. The organoids showed unhampered growth for up to ten days when directly cultured inside the ultra-thin FEP-foil cuvettes (**Figure 5a**). This verified that the FEP-foil properties facilitate sufficient gas exchange, and that this culturing technique ensures adequate nutrient and growth factor supply through the cuvette-opening. The matrix-embedded, delicate, mono-layered 3D cell cultures were fixed and DAPI-stained within the FEP-cuvettes ensuring the preservation of their morphology. Subsequent image acquisition resulted in high-quality 3D image stacks, which were fit for analysis with established quantitative image-based analysis tools. In particular, we applied our recently developed multiscale image analysis pipeline to the acquired image data ***(41)***. Based on the DAPI staining (**Figure 5b**), the cell nuclei were segmented (**Figure 5c**) and basic morphological features were extracted for individual organoids (see **Movie 5**). **Figure 5d** shows detailed views of segmented nuclei, indicating a good detection performance due to high image quality. Based on the nuclei segmentation data, surface areas and volumes of the organoids were approximated in 3D (**Figure 5e, Supplementary Table 2**). Furthermore, nuclei volume distributions within individual organoids were determined, which reveals differing median nuclei volumes for each organoid (**Figure 5f**).

## Discussion

The need to work with delicate three-dimensional (3D) cell cultures such as tissue sections, organoids and spheroids continues to rise. Hence, the demand for simple but versatile sample preparation methods for light sheet-based fluorescence microscopy (LSFM) has also increased considerably. In order to provide outstanding imaging properties, we developed ultra-thin FEP-foil chambers, in particular cuvettes, which are easily manufactured and adapted to a sample’s properties. We describe the fabrication of ultra-thin FEP-foil cuvettes as specimen holders for LSFM. We also demonstrate their applicability with representative 3D image stacks of 3D native and optically cleared whole organs, thick tissue sections, single organoids and dense organoid clusters.

In the pioneering days of light-sheet fluorescence microscopy, we introduced FEP-foil sample holders to study the 3D dynamic instability of 3D microtubule asters ***(11)***. These early FEP-foil holders were produced by wrapping a FEP-foil sheet around a glass capillary and gluing it along the capillary’s axis to form a cylindrical container. Although we obtained good image quality using FEP-cylinders, the technique held several drawbacks, including leakage and restricted rotation for multi-view imaging due to light scattering caused by the glue line. The ultra-thin FEP-foil cuvettes overcome such issues.

The production of ultra-thin FEP-cuvettes combines 3D printing and vacuum forming. Both technologies are affordable and easily available. They provide microscope developers and users with a lot of freedom for customisation. Using the newly developed flexible, resilient and seamless ultra-thin FEP-foil cuvettes as sample containers for LSFM simplifies sample preparation and handling of large specimens. In particular, the positioning of the specimen inside the cuvettes is much easier compared to rigid and fragile cuvette-like sample holders such as glass capillaries ***(15, 16)***. Another advantage of the new ultra-thin FEP-foil cuvettes is their compatibility with broad ranges of native and optically cleared biological specimens, mounting media and refractive indices. We verified a high tolerance against spherical aberration originating from refractive index mismatches with frequently used optical clearing solutions such as CUBIC2 (n=1.49) and ECi (n=1.58). This tolerance is based on the very thin wall of the cuvette (~12 µm), which results from stretching the heated FEP-foil (original thickness about 50 µm) over the positive moulds in the vacuum forming process. The foil stretching is given by the so-called *draw ratio* (=surface area/footprint). The calculated draw ratio, based on the geometry outlined in **Supplementary Figure 2d**, is 6. The measured draw ratio obtained from the average footprint and surface areas is around 4 (**Supplementary Figure 2d**, table). The draw ratio depends on several factors in the process of vacuum forming: the temperature of the foil, the intensity of negative pressure, the shape of the mould, the ratios of the mould’s geometry, the curvature of the edges and the mould’s material (**Supplementary Figure 2d**).

A further advantage is that both the shape and seamless nature of ultra-thin FEP-foil cuvettes allow for an unrestricted rotation of the sample for multiple-view imaging. This simplifies the 3D reconstruction of multiple-view data sets, which improves the image quality of highly scattering specimens. Furthermore, the images’ quality is suitable for the application of state-of-the-art multi-scale image analysis tools ***(41)*** and, thus, meets the requirements of quantitative methods for system-based analysis approaches ***(42)***.

Beyond what we demonstrated, ultra-thin FEP-foil cuvettes are also suitable for long-term live imaging, if temperature and gas exchange are controlled. FEP is an inert and completely bio-compatible material that does not interfere with the normal cell physiology ***(43)***. Since the cuvette’s walls are very thin, good exchange of O_2_ and CO_2_ is ensured. At the same time, the thin walls are impermeable to liquids. Thus, the composition of the internal medium is completely under control and constant, and no external contamination or dilution due to osmotic pressure can occur. Indeed, we cultured pancreatic organoids embedded in Matrigel inside the FEP-cuvette for over a week prior imaging. Moreover, we conducted live imaging to study the growth and morphogenesis of pancreatic organoids for over 120 hours inside a FEP-cuvette (*Till Moreth, Michael Koch, unpublished data*).

Besides FEP, other thermoplastic polymeric foils with a broad range of refractive indexes is available on the market (**Supplementary Table 3**). The production of specimen holders from these foils by vacuum forming will further extend the usefulness of this approach in light sheet fluorescence imaging.

In summary, ultra-thin FEP-foil cuvettes are “universal” sample holders that greatly simplify imaging of a broad spectrum of specimens ranging in size, from small to large, both native and optically cleared. Introducing increasingly user-friendly and efficient procedures for specimen mounting in LSFM will encourage even more life scientists to take advantage of light sheet microscopy to answer biological questions.

## Supporting information

Movie1_Ovary

Movie2_Brain

Movie3_PancreasOrganoids

Movie4_LiverOrganoids

Movie5_SegmentedOrganoid

SupplementaryMovie1_Kidney

## Acknowledgments

FP, EHKS, MK, LH, TM, KH thank the EU Horizon2020 project LSFM4LIFE (grant # 668350-2), the ZonMw-BMBF joint sponsored project “The Onconoid Hub” (grant # 114027003), and the DFG Cluster of Excellence Frankfurt “Macromolecular Complexes” (CEF-MCII) for funding. We thank Fabian Reinisch for helping with clearing and preparation of the murine kidney. We also thank Peter J. Verveer and Remko Dijkstra of Scientific Volume Imaging for support and suggestions for the reconstruction of multi-view data sets. We kindly thank Francesca Klinger (University of Tor Vergata, Rome, Italy) for providing GFP-c-kit murine ovaries, Meritxell Huch (Gurdon Institute, Cambridge, UK) and Lorenza Lazzari (Cell Factory Policlinico Milano, Milan, Italy) for cooperation in the culture of pancreas organoids (LSFM4LIFE project), and Luc van der Laan (Erasmus Medical Center, Rotterdam, Netherlands) for cooperation in the culture of liver organoids (“The Onconoid Hub” project).

## Authors’ contribution

KH performed both clearing and imaging of the murine hippocampus and of the murine ovary and measured and analysed the point spread function. MK cultured, isolated and imaged the human hepatic organoids. LH cultured, imaged and processed the pancreatic organoids. TM grew organoids in FEP cuvettes. MT isolated and stained the murine ovaries. EHKS modelled the effect of wall thickness and refractive index mismatch on the image quality. FP invented the ultra-thin FEP-foil cuvettes, designed and improved their fabrication process, fabricated the cuvettes, processed the LSFM data, supervised the work. FP and KH wrote the manuscript. All the authors contributed to writing the manuscript, revised and approved it.

## Supplementary Figures

**Supplementary Figure 1:**
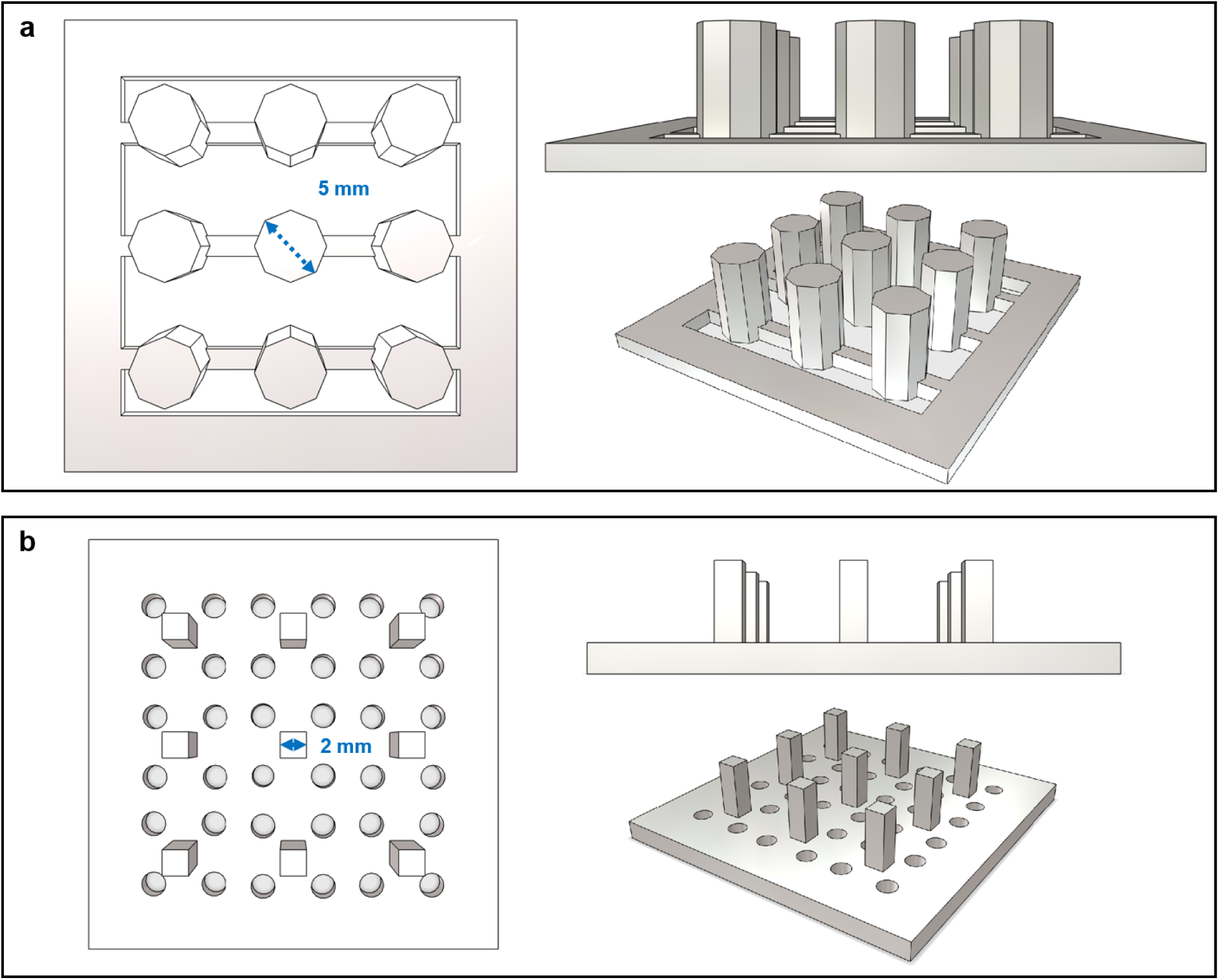
CAD-derived drawings of two positive moulds for ultra-thin FEP-foil cuvettes. Multiple cuvettes or other chambers are produced in parallel. Their number depends on their size and the extent of the base plate of the vacuum forming device. **(a)** Above, lateral and perspective views of the positive mould for ultra-thin FEP-foil cuvettes with an octagonal cross section and an diameter of 5 mm. **(b)** Comparable views of the positive mould for ultra-thin FEP-foil cuvettes with a square cross section and a width of 2 mm.

**Supplementary Figure 2:**
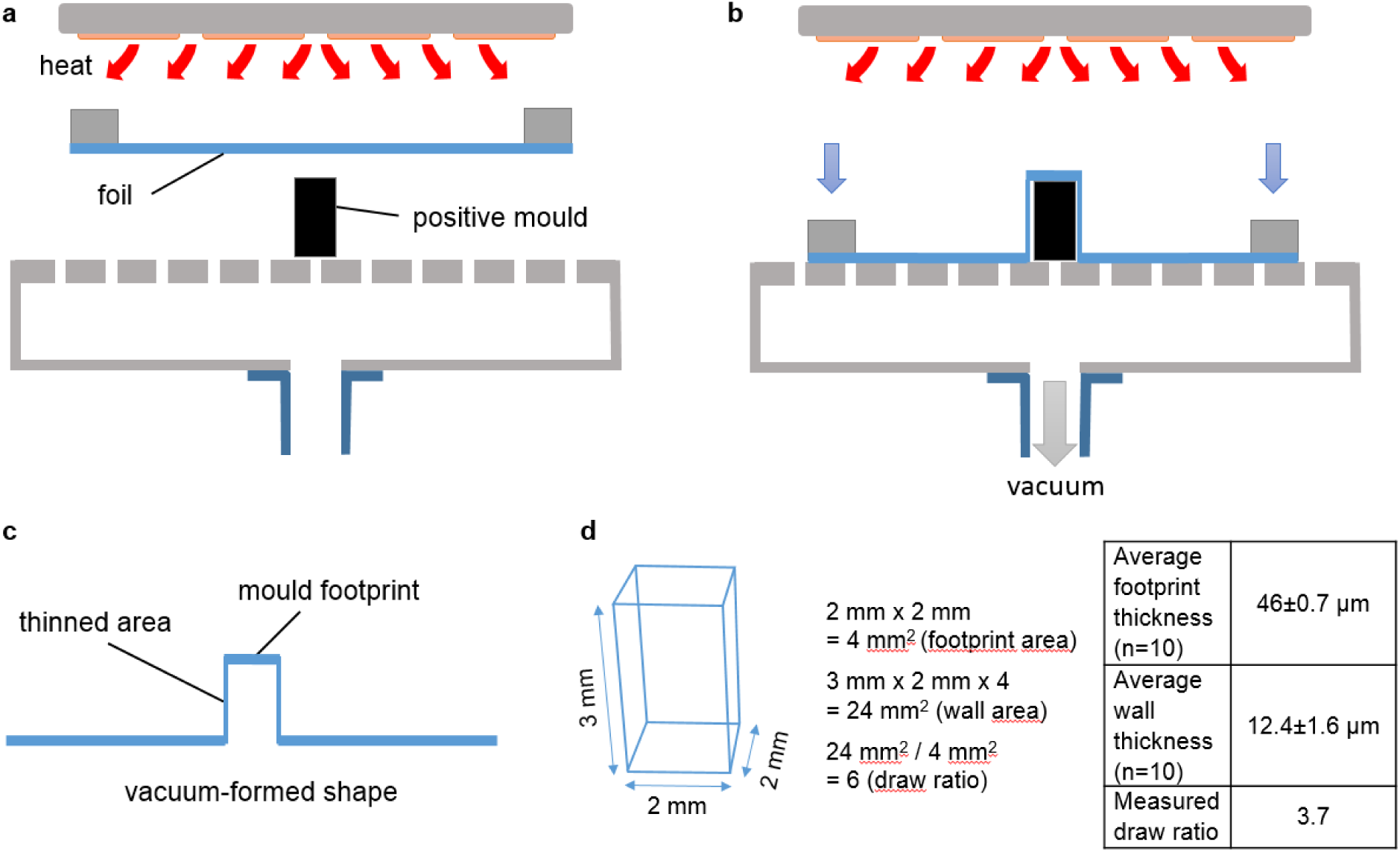
Vacuum forming process for ultra-thin FEP-foil cuvettes with a square cross section. **(a)** The thermoplastic FEP-foil is held firmly in a frame and heated to a target temperature close to its glass transition temperature. A positive mould is placed on the perforated vacuum plate. **(b)** Once the target temperature is achieved, the vacuum is established while the foil is stretched over the positive mould onto the perforated plate. **(c)** As soon as the foil has cooled down, the vacuum pressure is released and the ultra-thin FEP-foil cuvette is detached from the mould. **(d)** Calculation of the footprint area, the stretched wall area, and the draw ratio based on the geometry of a cuvette with a square cross section. The measured average footprint thickness and stretched wall thickness shown in the table result in an experimental draw ratio of about four.

**Supplementary Figure 3:**
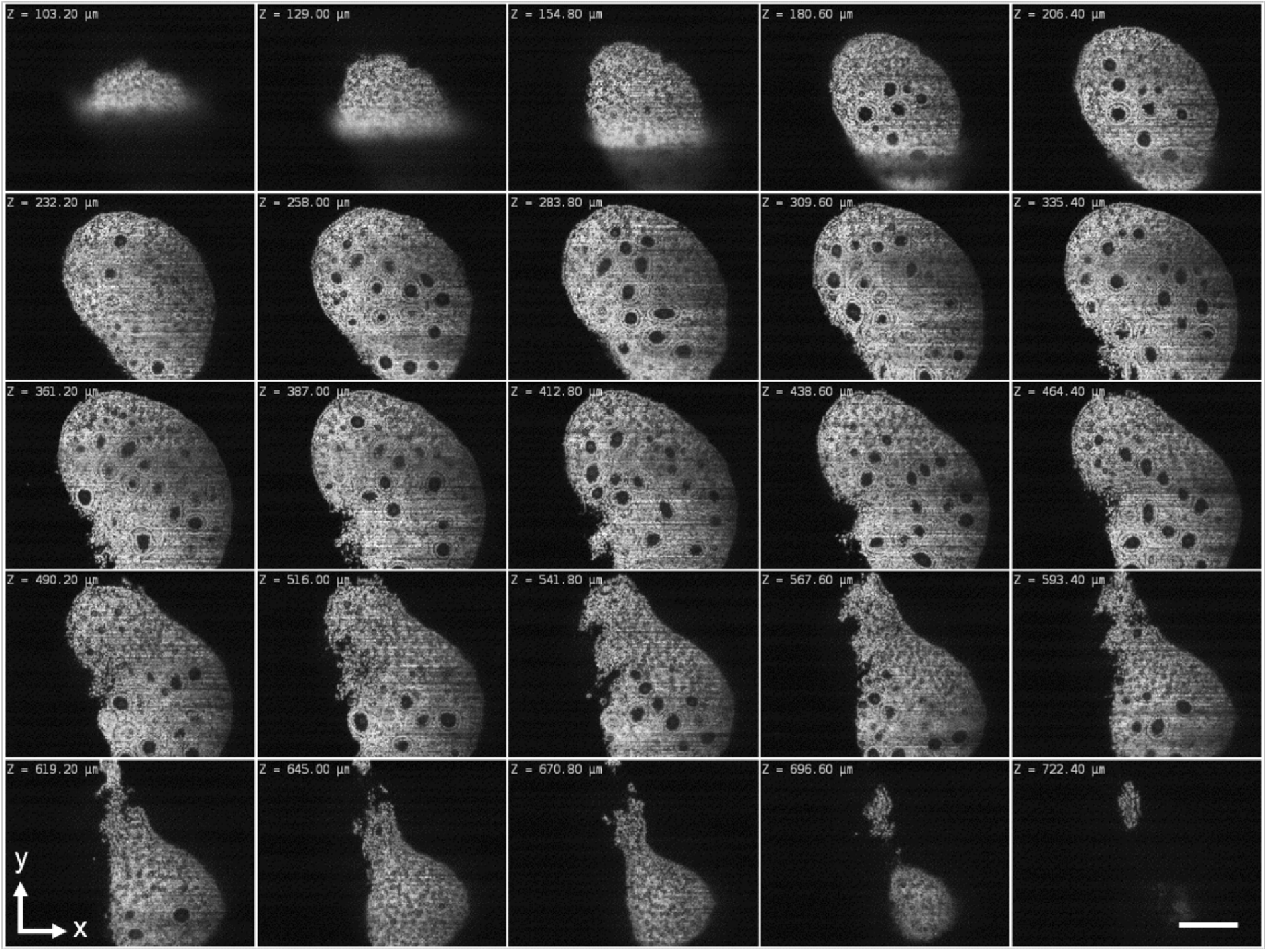
Selected frames from an image stack of a murine ovary optically cleared with CUBIC2. Clearing with CUBIC2 reduces the heterogeneity of the optical density, i.e. by decreasing scattering effects and hence blur, which allows the imaging of entire ovaries. In combination with ultra-thin FEP-foil cuvettes, a penetration depth of about 700 µm is easily achieved in LSFM. Staining: cell nuclei (DAPI). Microscope: mDSLM. Objective lenses: Epiplan-Neofluar 2.5x/0.06 (excitation). N-Achroplan 10x/0.3 (detection). Excitation wavelength: 405 nm. Bandpass detection filter: 447/25 nm. Scale bar: 300 µm.

**Supplementary Figure 4:**
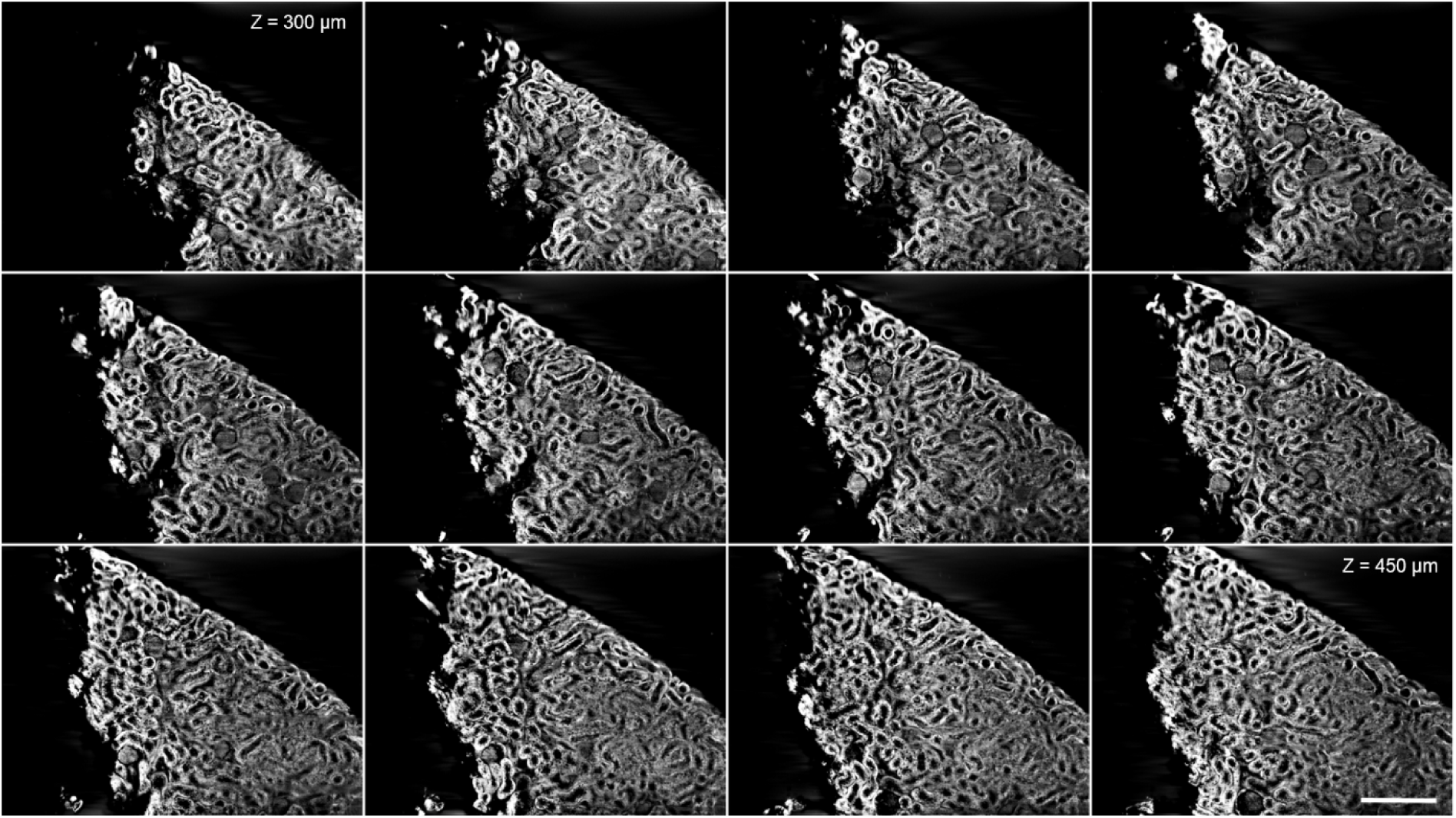
Selected frames from an image stack of a murine kidney section optically cleared with ethyl cinnamate (ECi). Images of an optically cleared murine kidney section in an ultra-thin FEP-foil cuvette. The autofluorescence signal of the kidney fragment was detected. The complex kidney ductal structure and the glomeruli are clearly visible in the images. Microscope: mDSLM. Objective lenses: Epiplan-Neofluar 2.5x/0.06 (excitation). N-Achroplan 10x/0.3 (detection). Excitation wavelength: 488 nm. Band pass detection filter: 525/50 nm. Scale bar: 200 µm.

**Supplementary Figure 5:**
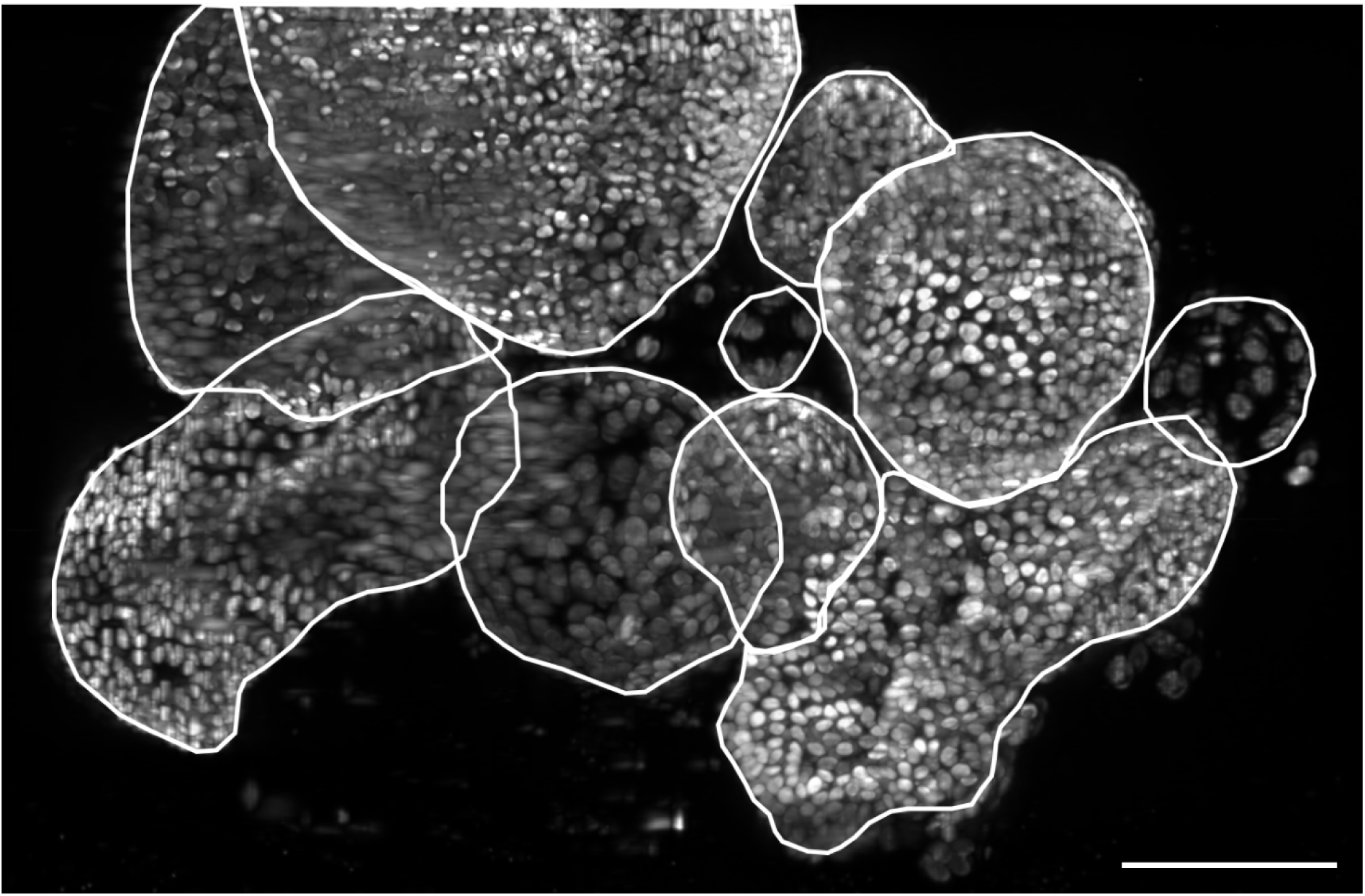
Maximum intensity XY-projection of a dense cluster of human hepatic organoids partially embedded in Matrigel. Maximum intensity XY-projection of a fused multi-view dataset (0, 90, 180, 270 degree) comprising 301 Z-planes. Individual organoids are highlighted with white contour lines. Staining: cell nuclei (DAPI). Microscope: mDSLM. Objective lenses: Epiplan-Neofluar 2.5x/0.06 (excitation). N-Achroplan 10x/0.3 (detection). Excitation wavelength: 405 nm. Bandpass detection filter: 447/25 nm. Scale bar: 100 µm.

**Supplementary Figure 6:**
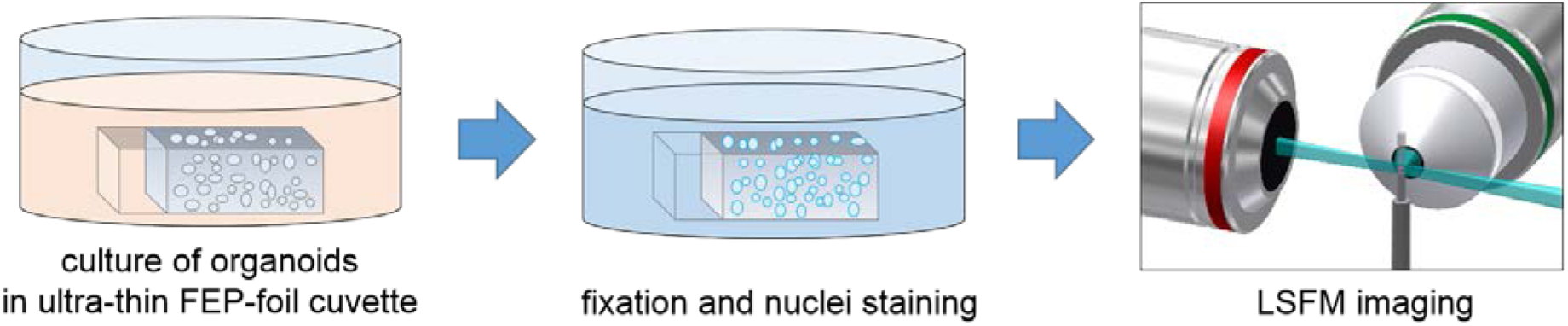
Long-term cultivation of pancreatic organoids inside ultra-thin FEP-foil cuvettes for LSFM. Murine pancreatic organoids were cultivated embedded in extracellular matrix Matrigel inside ultra-thin FEP-foil cuvettes. The FEP-cuvettes were placed in a 48-well plate, and fully covered with expansion medium to ensure growth factor and nutrient supply. The ultra-thin FEP-foil cuvettes enable optimal gas exchange and are suitable for a cultivation period of up to ten days. Since no sample transfers are required for fixation and staining, the organoid morphology is preserved. This is a prerequisite for high quality three-dimensional imaging.

## Supplementary tables

**Supplementary Table 1:** Raw data of measured Full Width Half Maximum (FWHM) values of Tetraspeck fluorescence beads excited with 488 nm imaged with an mDSLM with LSFM in PBS and CUBIC2. The expected values for the lateral and axial FWHMs are 1.8 µm and 14 µm respectively.

| | PBS (XY) [$\mu\text{m}$] | CUBIC2 (XY) [$\mu\text{m}$] | PBS (YZ) [$\mu\text{m}$] | CUBIC2 (YZ) [$\mu\text{m}$] |
| --- | --- | --- | --- | --- |
| **Bead 01** | 2.37 | 1.31 | 14.49 | 8.75 |
| **Bead 02** | 2.26 | 1.64 | 13.69 | 8.87 |
| **Bead 03** | 2.27 | 1.35 | 13.55 | 11.52 |
| **Bead 04** | 2.24 | 1.64 | 15.32 | 10.73 |
| **Bead 05** | 2.63 | 1.27 | 16.79 | 8.68 |
| **Bead 06** | 1.91 | 1.08 | 13.69 | 10.32 |
| **Bead 07** | 1.73 | 1.19 | 14.73 | 11.29 |
| **Bead 08** | 1.98 | 1.99 | 13.28 | 8.87 |
| **Bead 09** | 2.05 | 2.31 | 13.71 | 8.87 |
| **Bead 10** | 1.79 |  | 12.74 | 8.75 |
| **mean** | 2.15 | 1.5 | 14.2 | 9.78 |
| **standard deviation** | 0.26 | 0.379 | 1.12 | 1.11 |
| **Improvement** | $(2.15 \pm 0.26 - 1.5 \pm 0.38) / 2.15 \pm 0.26 = (30 \pm 22)\%$ | | $(14.2 \pm 1.12 - 9.8 \pm 1.11) / 14.2 \pm 1.12 = (31 \pm 11)\%$ | |

**Supplementary Table 2:** Summary of the morphology features drawn from the nuclei segmentation of the three different organoids displayed in Figure 5, pixel size: 0.654 x 0.654 µm², voxel size: 0.654 x 0.654 x 0.654 µm³.

|  | Organoid No. 1 | Organoid No. 2 | Organoid No. 3 |
| --- | --- | --- | --- |
| NucleiCount [#] | 823 | 803 | 261 |
| Volume [# of voxels] | 89605859 | 57111992 | 24797363 |
| SurfaceArea [# of pixels] | 993789 | 788656 | 423386 |
| MinCount [# of voxels] | 500 | 501 | 504 |
| MaxCount [# of voxels] | 16572 | 11959 | 14681 |
| MeanCount [# of voxels] | 4031 | 3252 | 3925 |
| StandardDeviationCount [# of voxel] | 2284 | 1848 | 2664 |

**Supplementary Table 3:** Thermoplastic polymer foils commercially available for vacuum-forming

| Material | Commercial name | Refractive index | Available foil thickness |
| --- | --- | --- | --- |
| Cyclo-olefin Polymer | ZEONEX, ZEONOR | 1.52 | 40µm-188µm |
| TetraFluorEthylene, Perfluorpropylene, FEP | LUMOX, Teflon | 1.34 | 12.5µm, 25µm, 50µm, 125µm |
| Polyamide | GRILAMID TR | 1.54 |  |
| Polyimide | KAPTON | 1.66 | from 7.9µm to 152µm |

## Movie captions

Movie 1: Three-dimensional surface rendering of an optically cleared murine ovary. Nuclei (grey) are stained with DAPI, oocytes (green) specifically express GFP-c-Kit. 360° rotation with 1° increments.

Movie 2: Three-dimensional maximum projection of an optically cleared hippocampus of a mouse expressing Thy1-GFP (green). 360° rotation with 10° increments.

Movie 3: Three-dimensional surface rendering of a native, immuno-fluorescence labelled human pancreatic organoid. DAPI-stained nuclei (grey), phalloidin-stained F-actin (green), immune-stained Sox9 (magenta).

Movie 4: Three-dimensional projection of a fused multi-view dataset of human hepatic organoids in Matrigel. DAPI-stained nuclei (grey), 360° rotation at 10° increments.

Movie 5: Three-dimensional surface rendering of nuclei-segmented murine pancreatic organoid. Colours represents individual detected nuclei.

